# No evidence for an S cone contribution to the human circadian response to light

**DOI:** 10.1101/763359

**Authors:** Manuel Spitschan, Rafael Lazar, Ebru Yetik, Christian Cajochen

**Affiliations:** Department of Experimental Psychology, University of Oxford, United Kingdom; Centre for Chronobiology, Psychiatric Hospital of the University of Basel (UPK), Switzerland; Transfaculty Research Platform Molecular and Cognitive Neurosciences, University of Basel, Switzerland

## Abstract

Exposure to even moderately bright, short-wavelength light in the evening can strongly suppress the production of melatonin and can delay our circadian rhythm. These effects are mediated by the retinohypothalamic pathway, connecting a subset of retinal ganglion cells to the circadian pacemaker in the suprachiasmatic nucleus (SCN) in the brain. These retinal ganglion cells directly express the photosensitive protein melanopsin, rendering them intrinsically photosensitive (ipRGCs). But ipRGCs also receive input from the classical photoreceptors — the cones and rods. Here, we examined whether the short-wavelength-sensitive (S) cones contribute to circadian photoreception by using lights which differed exclusively in the amount of S cone excitation by almost two orders of magnitude (ratio 1:83), but not in the excitation of long-wavelength-sensitive (L) and medium-wavelength-sensitive (M) cones, rods, and melanopsin. We find no evidence for a role of S cones in the acute alerting and melatonin supressing response to evening light exposure, pointing to an exclusive role of melanopsin in driving circadian responses.

---

To probe the role of S cones in circadian responses to light, we generated a pair of stimuli providing either minimal S cone stimulation, S–, or maximal S cone stimulation, S+ (Fig. 1a). The stimuli were designed to produce no differential stimulation of the L and M cones, the rods, and melanopsin (Fig. 1a, inset). We employed a spectrally tuneable light source consisting of ten different LED lights, which were individually adjustable in intensity, thereby producing complex mixtures of light which differed in the amount of S cone stimulation by a factor of ∼85, or equivalently, ∼1.92 units. The S cones play an important role in colour vision, encoding the blue-yellow dimension of colour vision. As a consequence, our S-cone isolating stimuli looked very different in colour (but not luminance, or ‘brightness’), with S– corresponding to an orangish, and S+ corresponding to a pinkish colour.

**Figure 1.**
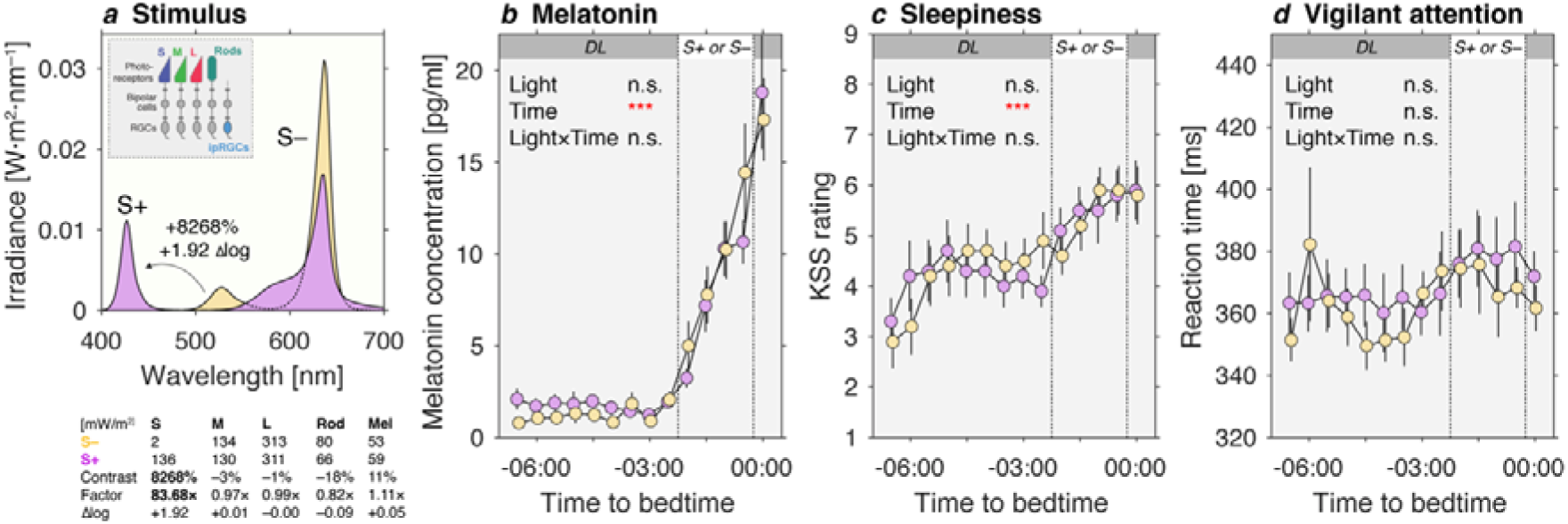
***a*** Spectral power distribution for the S-cone-isolating stimuli in peripheral presentation (annulus, inner ⍰ =11°, outer ⍰ = 58°), with minimal stimulation of L and M cones, rods, and melanopsin (S– = minimum S cone stimulation; S+ = maximum S cone stimulation, while retaining L and M cone, rod and melanopsin excitation). The difference in S cone stimulation can be specified as contrast (8268%), factor (83.68×), or log unit difference (+1.92 Δlog). Numbers are calculated from the actual spectrum measured in from the observer’s point of view. ***b*** Melatonin concentrations, with characteristic increase in melatonin in the evening. No differential effect is observed in the S– and S+ conditions. ***c*** Subjective sleepiness as measured using the Karolinska Sleepiness Scale (KSS), with characteristic increase in melatonin in the evening. No differential effect is observed in the S– and S+ conditions. ***d*** Vigilant attention, as measured using simple RT to an auditory beep. Data points with error bars are mean±1SEM. All statistical tests are described in the materials section.

With these stimuli in hand, we probed the human circadian timing system using melatonin suppression. Melatonin, which rises in concentration approximately two hours prior to habitual bedtime, can be strongly suppressed by short-wavelength light [1, 2]. In an in-laboratory within-subjected design under controlled lighting conditions, we found no difference in melatonin production when participants (n=10) were exposed to our two stimuli differing in S cone activation (Fig. 1b) from 150 to 30 minutes prior to their habitual bedtime. While a change in light stimulus by almost two orders of magnitude (1:100) is known to move the circadian response to light from no response to saturation [3], a change of that order of magnitude in only the S cones produced no difference in the production of evening melatonin. We also interrogated our stimuli affected subjective sleepiness (measured using the Karolinska Sleepiness Scale) and vigilant attention (measured using median reaction time to beeps, averaged over 50 trials). Neither sleepiness nor vigilant attention were modulated by S cones alone.

Our results further strengthen the notion that the most important modulator of circadian photoreception is melanopsin, and melanopsin only. In addition to our results ruling out the S cones as a driver of melatonin suppression, this emerging picture is supported by studies with people with red-green colour vision deficiencies affecting the L or M cones [4] and blind people with no cone-rod function at all [5, 6].

In the primate retina, some ipRGCs receive positive, excitatory input from the L and M cones and negative, inhibitory synaptic input from the S cones [7]. Previous research also exploiting the method of silent substitution found paradoxical responses of the pupil to flickering S- cone-isolating stimuli [8]. Our findings suggest that the circuit responsible for pupil control may recruit different ipRGCs than those involved in circadian photoreception [9, 10], or adapts to differences in cone input, or may have different temporal integration properties downstream.

Recent developments of lighting engineering and design have enabled the control of spectrum and intensity in the built environment. The lack of an S-cone mediated contribution to human circadian response to light is a key piece in the puzzle to optimising lighting for human health and well-being. In these considerations, the activation of melanopsin should be the only parameter when it comes to effects on the circadian system.

## Acknowledgements

M.S. is supported by a Sir Henry Wellcome Trust Fellowship (Wellcome Trust 204686/Z/16/Z) and a Junior Research Fellowship from Linare College, University of Oxford.

## Online Supplement: Materials and Methods

### Participant characteristics

Seventeen male participants (n=17) aged 18-35 were recruited to participate in the study (mean age±1SD: 24.1±2.72 years). Our participants were screened for sleep disruption (>5 on Pittsburgh Sleep Quality Index, PSQI, [1]), extreme morningness or eveningness (>27 or <11 on modified Horne & Östberg questionnaire, [2]), depressive symptoms (>27 on Center for Epidemiologic Studies Depression scale, CES-D, [3]), alcohol use disorder (>19 on Alcohol Use Disorders Identification Test, AUDIT, [4]), abnormal colour vision (assessed with Hardy-Rand-Rittler (HRR) plates), and visual acuity (at least 20/40 assessed using Snellen chart).

### Data exclusion and missing data

We had to exclude 7 of the 17 participants’ melatonin profiles due to low quality in the melatonin samples, due to contamination or insufficient saliva. Our analyses of the melatonin, sleepiness (KSS), and vigilant attention (median RT) data do not include these participants. Our analysis of the light questionnaire includes the full data from 17 participants. In some samples, the assays returned implausibly high melatonin samples (e.g. >30 pg/mL six hours before habitual bedtime), in which case we detected and removed outliers across participants but within-condition more than three scaled median absolute deviations (MAD) away from the median (implemented in MATLAB’s isoutlier function). Neither including participants with poor melatonin data quality, nor not removing the outliers rendered the statistical comparison (S– vs. S+) significant.

### Stimulus design and delivery

Visual stimuli were generated using a 10-primary LED-based light source (SpectraTune LAB, Ledmotive Technologies S.L, Barcelona, Spain) imaged onto a diffusing surface with independent 12-bit (4096 levels, including off) software control over the spectral emittance over each primary. Eight of the 10 primaries were relatively narrowband (427±16 nm [peak wavelength±FWHM at 100% intensity], CIE 1931 xy chromaticity: (0.17, 0.02); 445±20 nm, (0.17, 0.03); 465±24 nm (0.14, 0.07), 474±30 nm (0.13, 0.12), 504±31 nm (0.11, 0.58), 522±34 nm (0.19, 0.71), 636±19 nm (0.70, 0.30), 659±19 nm (0.72, 0.28). Two additional primaries were broadband LEDs: lime (558±120 nm, (0.43, 0.54)) and orange (596±83 nm, (0.57, 0.52)). A mask was placed in front of the diffusing surface, so as to provide an annular region with an outer diameter of 20 cm and an inner diameter of 3.5 cm, viewed at a viewing distance of 18 cm from a chin rest (annulus inner diameter: ∼11°, outer diameter ∼58°), thereby providing peripheral stimulation appropriate for circadian responses to light [5].

We generated our S-cone-selective stimuli using the method of silent substitution [6, 7]. In the method of silent substitution, pairs of spectra are generated as mixtures of the ten primaries lights which produce a difference in only one photoreceptor class (in this case, the *stimulated* S cones), while there is no difference in the other photoreceptors (in this case, the *silenced* L and M cones, rods, and melanopsin). This method has previously been used to examine the effect of melanopsin-only differences in lighting on melatonin suppression [8, 9] (but has a long history in vision science, see [7]).

To produce calibrated stimuli, we first measured the spectral radiance of each LED independently at 19 intensity levels (spaced at 5% increments from 5% to 100%, where 100% is maximum intensity) using a spectroradiometer (spectroval 1511, JETI Instruments GmbH, Jena, Germany). We addressed the typical changes in spectrum with increasing intensity by relying on an interpolation-based forward model our primaries (interpolating at unmeasured primary settings). Using this model, we generated two sets of settings for our primaries which would have the feature that they yielded maximum differential stimulation on the S cones, with minimal change in L and M cone, rod and melanopsin stimulation. These settings were simulatenously found using constrained minimisation routines implemented in MATLAB (`fmincon` SQP solver with global optimisation; 1000 trial points). In this procedure, we used the cone, rod and melanopsin spectral sensitivities [10] comprising the 10° Stockman-Sharpe cone fundamentals [11], the CIE V’(λ) function for the rods, and the standard curve for melanopsin [12]. Irradiance spectra measured in the corneal plane from the observer’s point of view are given in Table S1.

We achieved a stimulus with a difference of 8268% (factor 83.68×), or equivalently almost two log units (∼1.92 log difference), in S cone stimulation, with minimal stimulation of L and M cones, rods and melanopsin. The photopic illuminances were 168 lux for the S– condition (0.48, 0.26; pink appearance) and 173 lux for the S+ condition (0.61, 0.37; ‘pink’ appearance). The melanopic irradiance was 59 mW/m2 for the S+ condition and 53 mW/m2 for the S– condition.

Validating the spectra from this optimisation procedure, our stimuli demonstrated excellent silencing for the L and M cones (Fig. 1; –3% L cone contrast, –1% M cone contrast), and very good silencing for rods (–18%) and melanopsin (+11%), while providing an almost two log unit difference S cone stimulation. It is unlikely that these small nominal differences produce a meaningful physiological difference, given the very large and to our knowledge unparalleled difference in S cone stimulation.

### Protocol

The study took place in a dedicated light-, temperature- and humidity-controlled apartment comprising a double-room as well as a dedicated bathroom (see Appendix A in [13] for photograph). Upon arrival (30 minutes prior to protocol start), participants gave a urine sample for drug test (multi-drug panel test for AMP, BZD, COC, MOR/OPI, MTD and THC; exclusion if positive; nal von minden, Den Haag, Netherlands) and accommodated to the laboratory. Then, the protocol began, lasting from 6.5 hours before habitual bedtime to habitual time. Every 30 minutes, participants completed an alertness assessment using simple auditory reaction time task, the Karolinska Sleepiness Scale (KSS), and gave a saliva sample using Salivettes in dim light provided by room illumination (photopic illuminance in the corneal plane <8 lux). From 2.5 hours to 0.5 hours prior to the habitual bedtime, participants were either exposed to the S– or S+ stimuli in 20 minute sections, yielding a total of 80 minutes of light exposure to the experimental stimulus.

Fixation and eye opening were verified using a video-based head-mounted eye tracker (Pupil Labs GmbH, Berlin, Germany). Participants had access to water throughout the experiment but no food or other drinks. Participants were allowed to spend their time reading, studying, playing Nintendo GameBoy (illuminance at cornea <8 lux), or other activities no involving additional light exposure. Smartphones and other electronic devices were removed from the experiment suite. All experiments took place between November 2018 and June 2019. All sessions took place one week from another and condition order was randomised between participants. From one week prior to the experiment to the second session, participants were instructed to adhere to regular bedtimes (±30 minutes) and wore actigraphy devices (Condor Instruments, São Paolo, Brasil). On the day of the experiment, participants were asked to refrain from caffeine consumption after noon.

### Salivary melatonin

Saliva samples (at least 1 mL) were collected at 30-minute intervals using Salivettes (Sarstedt AG, Sevelen, Switzerland), which were immediately centrifuged and frozen at −20° for later assay. Melatonin was measured using a direct double-antibody radioimmunoassay (analytical sensitivity 0.2 pg/mL, functional minimum detectable dose of 0.65 pg/ml; Bühlmann Laboratories AG, Allschwil, Switzerland).

### Vigilant Attention

Vigilant Attention was measured using a custom-made simple auditory reaction time task programmed in Psychtoolbox and MATLAB (The Mathworks, Natick, MA). Participants were presented with a tone emitted from a loudspeaker and were instructed to press as quickly as possible to the tone using a Playstation-like gamepad. ISI was randomly set to 5-8 seconds. Median reaction times were calculated from 50 trials.

### In-laboratory light questionnaire

Participants were asked to rate or respond to various aspects of the light exposure using a 6-question, 7-item Likert scale questionnaire. This questionnaire was administered in German. The questions were about the *comfort of light* (“Allgemein ist das Licht angenehm”; überhaupt nicht [1] – sehr stark [7]), the *perceived brightness* (“Wie empfinden Sie die Helligkeit des Lichtes?”; sehr dunkel [1] – sehr hell [7]), *light level preference* (“Ich hätte es lieber…”; deutlich dunkler [1] – deutlich heller [7]), *glare* (“Dieses Licht blendet mich”; überhaupt nicht [1] – sehr stark [7]), *colour temperature* (“Wie empfinden Sie die Lichtfarbe?”; sehr kalt [1] – sehr warm [7]) and *general well-being* (“Wie fühlen Sie sich im Moment?”; unwohl [1] – wohl [7]).

### Karolinska Sleepiness Scale (KSS)

We used the *German* version of the Karolinska Sleepiness Scale (“Bitte bewerten Sie Ihre Müdigkeit” (“sehr wach” [1], “wach” [3], “weder wach noch müde” [5], “müde, aber keine Probleme, wach zu bleiben” [7], “sehr müde, große Probleme, wach zu bleiben, mit dem Schlaf kämpfend” [9]).

### Statistical analysis

We modelled our data using a linear mixed-effects model, modelling subjects as a random-effects, and condition (S+ or S–) and sample number (with sample #14 corresponding to habitual bedtime) as fixed effects, along with the interaction between condition and sample. In Wilkinson-Rogers notation, our model is specified as

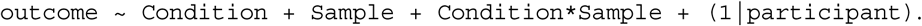

We subjected the melatonin concentrations (given in pg/mL), the median reaction times (given in seconds), the KSS scores, and the 7-item Likert scale questions from our light questionnaire to this model implemented in lme4 [14], with further statistical tests performed using lmerTest [15]. All F values reported reflect a Type III ANOVA, implementing Satterthwaite’s method for determining the degrees of freedom.

For melatonin, we found a significant effect of sample time (F(1, 95.607) = 20.7347, p = 1.554e-05 [***]), but no differences between conditions (F(1, 94.098) = 0.9073, p = 0.3433), and no interaction (F(1, 93.994) = 1.4463, p = 0.2321). The effect of sample time on melatonin is well evident in Fig. 1b, indicating the increase of melatonin with approaching habitual bedtime.

For sleepiness (KSS), we found an effect of sample time (F(1, 116) = 15.9744, p = 0.0001132 [***]), but no differences across conditions (F(1, 116) = 0.8745, p = 0.3516608), and no interaction (F(1, 116) = 0.8844, p = 0.3489551). The effect of sample time is also evident in Fig. 1c, where we see that the sleepiness increases, the closer the sample time is to the participant’s habitual bedtime. For alertness (median reaction time), we found no effect of sample time (F(1, 116) = 0.3381, p =0.56210, no differences across conditions (F(1, 116) = 0.7979, p = 0.3736), and no interaction between condition and time (F(1, 116) = 0.7391, p = 0.3917).

For the lighting questionnaire, which we sampled at the same intervals as the saliva samples and the KSS, we only compared the ratings of acute stimulus (delivered 02:30 to 00:30h before habitual bedtime). In this analysis, there were no significant differences between the two conditions, or across samples for the comfort of the light (condition: F(1, 116) = 0.8120, p = 0.3694; sample: F(1, 116) = 1.9172, p = 0.1688; condition x sample: F(1, 116) = 0.5559, p = 0.4574), the light level preference (condition: F(1, 116) = 0.5132, p = 0.4752; sample: F(1, 116) = 2.7313, p = 0.1011; condition x sample: F(1, 116) = 0.2586, p = 0.6121), or general well-being (condition: F(1, 115.06) = 0.0656, p = 0.7983; sample: F(1, 115.06) = 0.4380, p = 0.5094; condition x sample: F(1, 115.06) = 0.0216, p = 0.8834). Both perceived brightness (condition: F(1, 116) = 0.4403, p = 0.508294; sample: F(1, 116) = 10.8308, p = 0.001323 [**]; condition x sample: F(1, 116) = 0.5522, p = 0.458904) and glare (condition: F(1, 116) = 2.5974, p = 0.10976; sample: F(1, 116) = 6.3895, p = 0.01282 [*]; condition x sample: F(1, 116) = 2.1307, p = 0.14708) showed an effect of sample, but not of condition, suggesting some adaptation to the acute stimulus light in terms of brightness and glare perception. Critically, we find a significant effect of the two conditions on perceived colour temperature (F(1, 116) = 4.9736, p = 0.02766 [*]), along with time-dependent effect (F(1, 116) = 3.9827, p = 0.04831 [*]), but no interaction (F(1, 116) = 2.4344, p = 0.12142). The difference in perceived colour temperature is expected due to the different colour appearance of the two stimulus conditions. The rating data are shown in Fig. S1.

### Ethical approval

This study was approved by the cantonal ethics commission (Ethikkommission Nordwest- und Zentralschweiz, PB_2018-00164 – 280/90) and was conducted in accordance with the Swiss law and according to the Declaration of Helsinki.

## Online Supplement: Figures

**Figure S1.**
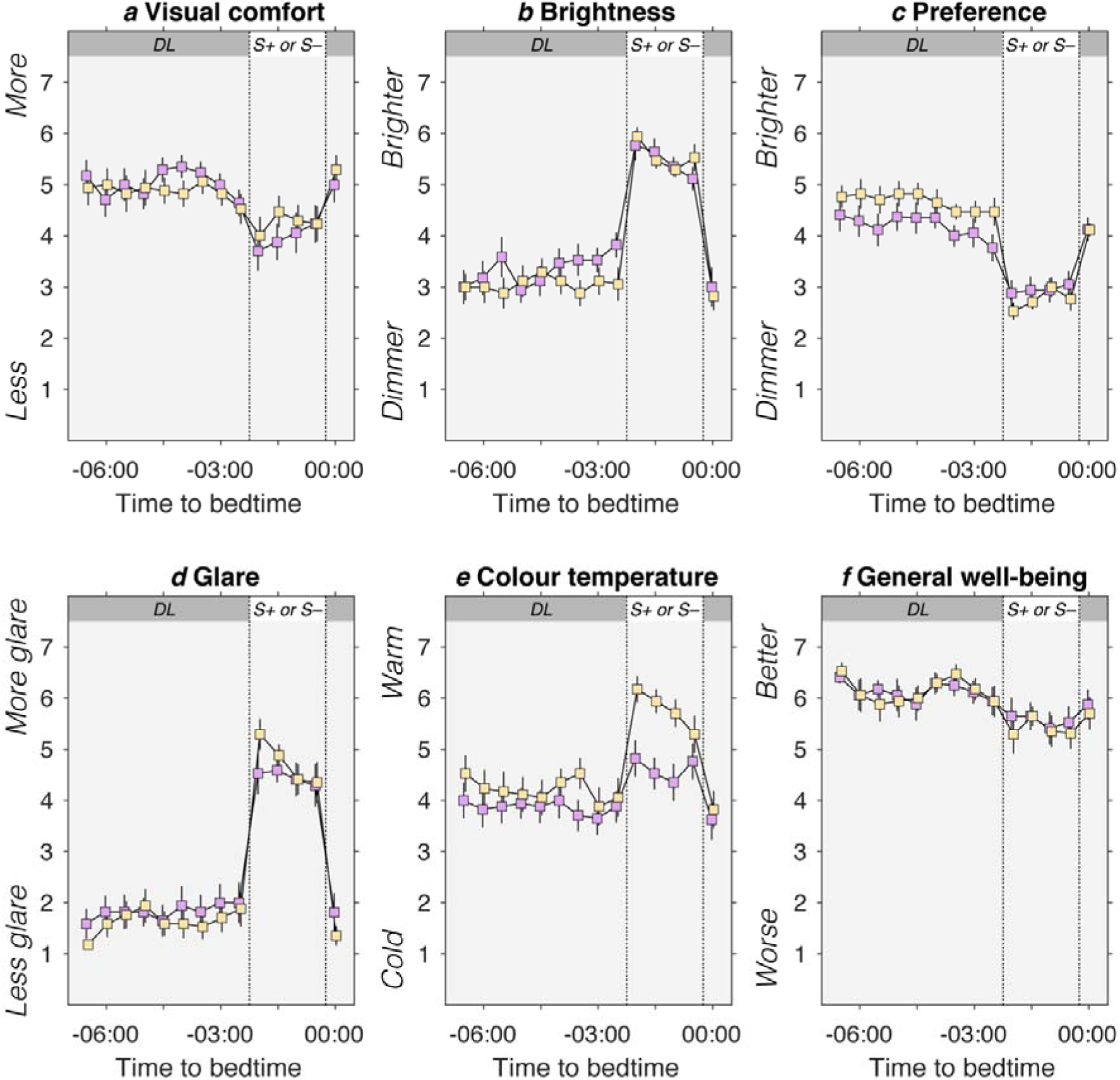
Rating results for visual comfort (***a***), brightness (***b***), light level preference (***c***), glare (***d***), colour temperature (***e***), and general well-being (***f***). Details about questions are given in the text.

